# Circulating Immune Profiling Reveals Immune Signatures Associated with Disease Stage and Outcome in Endometrial Cancer

**DOI:** 10.64898/2026.07.21.739790

**Authors:** Raquel Piñeiro-Pérez, Ana Vilar, Efigenia Arias, Victoria Sampayo, Alicia Abalo-Piñeiro, Carmela Rodríguez, Alexandra Cortegoso, Raúl Márquez, Eva Díaz, Gema Moreno-Bueno, Isabel Palacio, Sonia Blanco-Prieto, Lidia Vázquez-Tuñas, Isaura Fernández-Pérez, Ester Munera-Maravilla, Silvia Calabuig, Cristina Caballero, Ana Herrero, Rafael López-López, Juan Cueva, Juan E. Viñuela-Roldán, Laura Muinelo-Romay

**Affiliations:** Translational Medical Oncology (Oncomet)/ Liquid Biopsy Analysis Unit, Health Research Institute of Santiago (IDIS), Complexo Hospitalario Universitario de Santiago de Compostela, 15706, Santiago de Compostela, Spain; Universidade de Santiago de Compostela (USC), Santiago de Compostela, Spain; Galician Precision Oncology Research Group (ONCOGAL) Medicine and Dentistry School, Universidade de Santiago de Compostela, Santiago de Compostela, Spain; Gynecology Department, University Clinical Hospital of Santiago de Compostela (SERGAS), Santiago de Compostela, Spain; Medical Oncology Department, MD Anderson Cancer Center, C/Arturo Soria 270, 28033, Madrid, Spain; Fundación MD Anderson Internacional, Madrid, Spain; Centro de Investigación Biomédica en Red de Cáncer (CIBERONC) Monforte de Lemos 3, 5, 28029 Madrid, Spain; Hospital Universitario Central de Asturias, Department of Medical Oncology, Oviedo, Spain; Instituto de Investigación Sanitaria Galicia Sur, Vigo, Spain; Medical Oncology Department, Hospital Álvaro Cunqueiro-Complejo Hospitalario Universitario de Vigo, Vigo, Spain; Fundación para la Investigación del Hospital General Universitario de Valencia, Valencia, Spain; Pathology Department, Universitat de València, Valencia, Spain; Servicio de Oncología Médica, Hospital General Universitario de Valencia, Valencia, Spain; Hospital Universitario Miguel Servet, Zaragoza, Spain; Servicio de Oncología Médica, Hospital Universitario Miguel Servet, Zaragoza, Spain; Inmunología, Servicio de Análisis Clínicos, Complexo Hospitalario Universitario de Santiago de Compostela, SERGAS, A Coruña, Spain

**Keywords:** Endometrial cancer, predictive biomarkers, tumor immunology, flow cytometry, liquid biopsy

## Abstract

Although the tumor immune microenvironment has been studied in endometrial cancer, the systemic immune alterations associated with disease progression and their potential prognostic significance remain poorly defined. In this study, peripheral blood immune subsets were characterized by multiparametric flow cytometry in 67 patients with EC and 20 healthy controls, including dendritic cells, MDSCs, T-cell subsets, NK cells, and exhaustion and senescence associated markers. Immune profiles were similar between healthy controls and patients with early-stage disease, whereas advanced tumors showed marked changes, including dendritic cell expansion, reduced CD4^+^ T-cell proportions, and increased frequencies of CD8^+^CD27^-^CD28^-^ and CD8^+^CD57^+^ populations. Among clinicopathologic features, MDSC levels were associated with tumor grade and myometrial infiltration, while regulatory T cells were increased in TP53-mutated and microsatellite-stable tumors. In the advanced cohort (n=31), non-responders frequently displayed elevated CD8^+^, CD8^+^CD27^-^CD28^-^, and CD8^+^CD57^+^ levels alongside decreased CD27^+^CD28^+^ proportions. In multivariable Cox models, higher baseline CD8^+^ (HR 1.13), CD8^+^ CD27^-^CD28^-^ (HR 1.04), and CD8^+^CD57^+^ proportions (HR 1.07; all p<0.05) were independently associated with shorter PFS, whereas higher CD27^+^CD28^+^ levels were associated with improved PFS. CD27^-^CD28^-^ proportions were also linked to worse PFS by Kaplan-Meier analysis (HR 5.1, log-rank p=0.005). Longitudinal analysis (n=28) showed that senescent-like lymphocyte levels remained associated with progression across timepoints (OR 18.80), whereas total CD8^+^ proportions diverged progressively, reaching significance only from 6 months onward. Together, these findings identify systemic immune remodeling as a characteristic of advanced endometrial cancer and support the potential of circulating immune profiling for patient stratification and prognostic assessment, pending validation in larger prospective cohorts.

## Introduction

Endometrial cancer (EC) ranks as the fourth most diagnosed cancer in women and the eighth leading cause of cancer-related deaths among females globally [1]. While early-stage EC is often effectively treated with surgery and adjuvant therapies, patients with advanced-stage disease face a poorer prognosis and fewer treatment options. Advanced EC includes stage III or IV disease, characterized by the spread of cancer beyond the uterus and cervix to nearby organs or distant sites [2]. For many years, carboplatin plus paclitaxel was considered the standard first-line treatment for advanced EC. However, this therapeutic landscape has evolved substantially in recent years following the introduction of a molecular classification of EC, which has improved tumor stratification and revealed distinct biological subtypes with different clinical behaviors and therapeutic vulnerabilities [3, 4]. This stratification has paved the way for the development and clinical implementation of targeted therapies, including immunotherapy.

In this context, immune checkpoint inhibitors, such as pembrolizumab and nivolumab, have emerged as effective therapeutic options, particularly for tumors exhibiting microsatellite instability-high (MSI-H) or mismatch repair deficiency (dMMR), which are characterized by high tumor mutational burden and increased immunogenicity [5, 6, 7].

Taking into account this therapeutic context, the identification of accurate predictive biomarkers plays a crucial role in EC management. While clinical trials are exploring novel targeted therapies, implementation in practice is hindered by a lack of reliable and dynamic predictive tools [8]. Therefore, the study of circulating biomarkers has gained high interest as an alternative to personalize therapies while accounting for tumor heterogeneity and the dynamics of tumor cells and the microenvironment [9, 10].

Several studies have highlighted the relevance of immune microenvironment for predicting the tumor aggressiveness and the therapy response. The presence of tumor-infiltrating lymphocytes (TILs), particularly cytotoxic CD8^+^ T cells, has been associated with improved disease-free survival (DFS) and overall survival (OS) in EC [5, 11]. Most of these studies are focused on the primary tumor microenvironment without assessing the patient’s systemic immune status, which is linked to tumor development, prognosis, and treatment response [12, 13]. Beyond local immune changes within the tumor microenvironment, EC may also induce systemic immune alterations that are reflected in the peripheral blood immune compartment. In fact, systemic inflammatory markers such as the neutrophil-to-lymphocyte ratio (NLR) have been investigated for their prognostic value, with elevated NLR linked to a more aggressive evolution [14]. More recently, levels of CD8^+^CD28^-^ and CD8^+^PD1^+^ T cells have been associated with high-risk EC [15]. However, there is a lack of a detailed phenotypic characterization of different immune cell populations in the peripheral blood of these patients to define its utility for predicting treatment response and monitoring disease progression.

Building on this rationale, the present study provides a comprehensive characterization of circulating immune cell populations in patients with EC. By analyzing peripheral blood samples, we sought to identify alterations in the proportions of specific immune cell subsets and explore their associations with clinicopathologic features and clinical outcomes. Through this approach, we observed immune signatures in advanced endometrial cancer suggestive of increased suppressive mechanisms, involving both regulatory cells and immune checkpoint-associated phenotypes. Additionally, by tracking the longitudinal evolution of these immune populations during treatment, we assessed how temporal changes in immune cell composition were associated with therapeutic response and disease progression. Collectively, these findings provide a rationale for further investigating circulating lymphocyte subsets as exploratory biomarkers in EC.

## MATERIALS AND METHODS

### Patient Recruitment

Patients and healthy people were enrolled in the study between September 2022 and September 2024. Patients with localized endometrial tumors were recruited at the Gynecologic Department from the University Hospital of Santiago de Compostela prior to the surgical intervention. Patients with advanced EC were recruited at the Oncology Departments from the University Hospital of Santiago de Compostela, Valencia General Hospital, Miguel Servet University Hospital, MD Anderson Cancer Center, and Álvaro Cunqueiro University Hospital.

Eligible patients were aged 18 years or older, had a confirmed histopathological diagnosis of EC of any histological subtype (excluding sarcomas), and, in the case of advanced disease, patients had to be candidates to receive one of the following treatment regimens: first-line chemotherapy (with carboplatin-paclitaxel or doxorubicin), first-line hormonal therapy, or first-, second- or third-line immunotherapy with anti-PD1 treatment. Patients with any concomitant malignant tumor at the time of enrollment or diagnosed within the previous five years were excluded. Blood samples were collected from 67 (aged 69±10.59 years) patients diagnosed with EC spanning FIGO stages I to IV and grades 1 to 3. Clinical characteristics of the cohort are detailed in Table 1. In addition, 20 healthy women (aged 63±5.43 years) were also invited to participate in the study as a control cohort.

**Table 1.** Clinical variables of patients with localized and advanced EC. Summary of clinical characteristics, including histological subtype, tumor grade, FIGO stage, disease progression status, depth of myometrial invasion, and death status.

| CLINICAL VARIABLES | N (%) |  | N (%) |
| --- | --- | --- | --- |
| **Histology** |  | **Grade** |  |
| EEC | 49 (73,13%) | G1 | 25 (37,31%) |
| USC | 12 (17,91%) | G2 | 16 (23,88%) |
| Clear Cell | 3 (4,48%) | G3 | 24 (35,82%) |
| Neuroendocrine | 1 (1,49%) | Unknown | 2 (2,99%) |
| Unknown | 2 (2,99%) |  |  |
| **FIGO Stage** |  | **Progression Disease** |  |
| I | 27 (40,3%) | Yes | 14 (20,89%) |
| II | 3 (4,48%) | No | 53 (79,10%) |
| III | 3 (4,48%) |  |  |
| IV | 30 (44,78%) |  |  |
| Unknown | 4 (5,97%) |  |  |
| **Myometrial infiltration** |  | **Exitus** |  |
| No | 28 (41,79%) | Yes | 8 (11,9%) |
| Yes | 33 (49,25%) | No | 59 (88,1%) |
| Unknown | 6 (8,96%) |  |  |

### Sample collection and processing

Peripheral blood samples were collected at baseline, prior to initiation of therapy, in 3 mL EDTA tubes supplemented with Transfix® reagent (100 µL/mL, TFB-20-01). Samples were stored at 4 °C and analyzed within seven days. Whole blood (120 µL per tube) was divided into seven tubes, each containing a distinct antibody panel (29 antibodies in total). Detailed antibody panels are provided in Supplementary Table 1. Samples were incubated with antibodies for 30 minutes at room temperature in the dark, followed by red blood cell lysis with FACS Lysing solution (BD Biosciences). After centrifugation (1,500 rpm, 7 minutes, brake on), cells were washed and resuspended in 1 mL FACS Flow buffer for cytometric acquisition.

### Flow Cytometry Acquisition and Analysis

Samples were analyzed in a FACSCanto II cytometer (BD Biosciences, San Jose, CA, USA) using FACSDiva software. Gates and thresholds were set based on fluorescence intensity, light scatter characteristics (forward and side scatter), and the exclusion of dead cells and cellular debris, ensuring accurate identification of immune cells. Analysis was performed using FlowJo v10 (Tree Star Inc., Ashland, OR, USA).

### Immune Phenotyping and Gating Strategy

Immune cell subsets were identified using a standardized sequential gating strategy on singlet, viable leukocytes. After exclusion of doublets (FSC-A vs FSC-H) and debris based on forward and side scatter, leukocytes were gated on the lymphocyte or myeloid regions according to scatter properties. Dendritic cells were defined as lineage-negative (CD3^-^CD14^-^CD19^-^CD56^-^ FITC) HLA-DR^+^ (DR PerCP) cells and further subdivided into plasmacytoid dendritic cells (CD123 PE) and myeloid dendritic cells (CD11c APC). Myeloid-derived suppressor cells (MDSCs) were identified as HLA-DR^low/^-^ (DR PerCP^low/^-^) CD11b PE^+^ CD33 APC^+^ cells. Regulatory T cells (Tregs) were identified within the CD4^+^ T-cell compartment using CD25 and CD127 expression, and were defined as CD4^+^CD25^+^CD127low/^-^ cells. T-cell subsets were characterized in the CD45^+^ PerCP CD3^+^ FITC population, with CD4^+^ APC and CD8^+^ PE distinguishing CD4^+^ and CD8^+^ T cells. NK cells were defined as CD3^-^ PerCP CD56^+^ PE lymphocytes. Senescence and exhaustion associated T cell subsets were evaluated using dedicated panels including CD3 V450, CD8 APC, CD27 PerCP, CD28 PE, CD57 FITC/PerCP, PD-1/CD279 PE-Cy7 and KLRG1 APC-Cy7.

Within CD8^+^ T cells, mature cytotoxic cells were defined as CD27^+^CD28^+^, whereas late-differentiated CD27^-^ CD28^-^ cells were considered senescent-like. Senescent-like cells were also identified as CD57^+^ and/or KLRG1^+^CD8^+^ T cells, and PD-1/CD279 expression was additionally quantified within these subsets as an exhaustion associated marker.

Fluorescence compensation matrices were generated from single-stained study samples for each fluorochrome, and positivity thresholds for PD-1, KLRG1 and CD57 were set based on fluorescence intensity distributions and clearly negative internal populations within each panel; once established, gates were kept constant across all samples.

### Statistical Analysis

Statistical analyses were performed using GraphPad Prism software (version 10.11, GraphPad Software, San Diego, CA, USA) and R (version 4.4, R Foundation for Statistical Computing, Vienna, Austria). Immune cell proportions were expressed as percentages and analyzed as continuous variables unless otherwise specified. Because immune cell distributions were non-Gaussian, non-parametric tests were used throughout. Comparisons between two independent groups were performed using the Mann-Whitney U test. Comparisons among more than two groups were performed using the Kruskal-Wallis test followed by Dunn’s multiple-comparisons test. Adjusted *p* values from Dunn’s test were reported where applicable to control for multiple testing.

For exploratory clinical association analyses, immune variables were summarized and compared across clinical groups using the *gtsummary* package in R. For selected categorical analyses and Kaplan-Meier plots, immune cell variables were dichotomized at the cohort median into high and low groups. Benjamini-Hochberg false discovery rate (FDR) correction was additionally applied to the p-values within each clinicopathologic variable, and the resulting q-values are reported alongside nominal p-values. A binary composite CD8^+^ phenotype was defined as low mature and high senescent-like status, coded as 1 for patients with low mature CD8^+^CD27^+^CD28^+^ proportions and high senescent-like CD8^+^ proportions, and as 0 for all other combinations.

Progression-free survival (PFS) was calculated from the start date of first-line treatment to radiological progression confirmed by CT scan. Kaplan-Meier curves were generated in RStudio, and differences between groups were assessed using log-rank tests. Univariate Cox proportional hazards models were fitted to evaluate the association between baseline immune cell proportions and PFS. For exploratory multivariable analyses, Cox models were restricted for selected immune variables and were adjusted for age, histology, grade, and myometrial infiltration. Hazard ratios (HRs) with 95% confidence intervals (CIs) were reported per 1% increase in immune cell proportion. The proportional hazards assumption was assessed using Schoenfeld residuals in the final models.

Spearman’s rank correlation analysis was performed to evaluate relationships between immune cell subsets in the advanced cohort.

For longitudinal categorical analyses, immune variables were compared across timepoints and their association with disease progression was assessed using Fisher’s exact test when appropriate. Odds ratios (ORs) with 95% confidence intervals (CIs) were calculated. In exploratory analyses, logistic regression models were used to estimate the association between immune cell levels and progression status over time.

## RESULTS

### Differential Immune Cell Profiles in Healthy Controls, Early-Stage and Advanced EC

A total of 87 individuals were included for the study, comprising 67 patients with EC (32 with localized disease and 35 with advanced or recurrent disease, including stages IIIC to IVB) and 20 age-matched healthy controls. Patient characteristics are shown in Table 1. Peripheral blood samples were analyzed by flow cytometry to profile immune cell subsets across myeloid and lymphoid compartments (Supplementary Figures 1 and 2). Key populations examined included dendritic cells, MDSCs, CD3^+^, CD4^+^ and CD8^+^ T cells, among others.

We first focused on the dendritic cells, known for their role in antigen presentation and activation of T cells, and defined as lineage negative (CD19^-^, CD3^-^, CD14^-^ and CD56^-^), positive for HLA-DR and for CD123 (Plasmacytoid) or CD11c (Myeloid). Dendritic-cell proportions were higher in advanced EC than in control and localized EC groups, with a significant overall group effect (Kruskal-Wallis global *p* = 0.027) (Figure 1.A). No significant differences were observed between controls and localized patients, suggesting that dendritic cell expansion may correlate with disease dissemination rather than early tumor presence.

**Figure 1.**
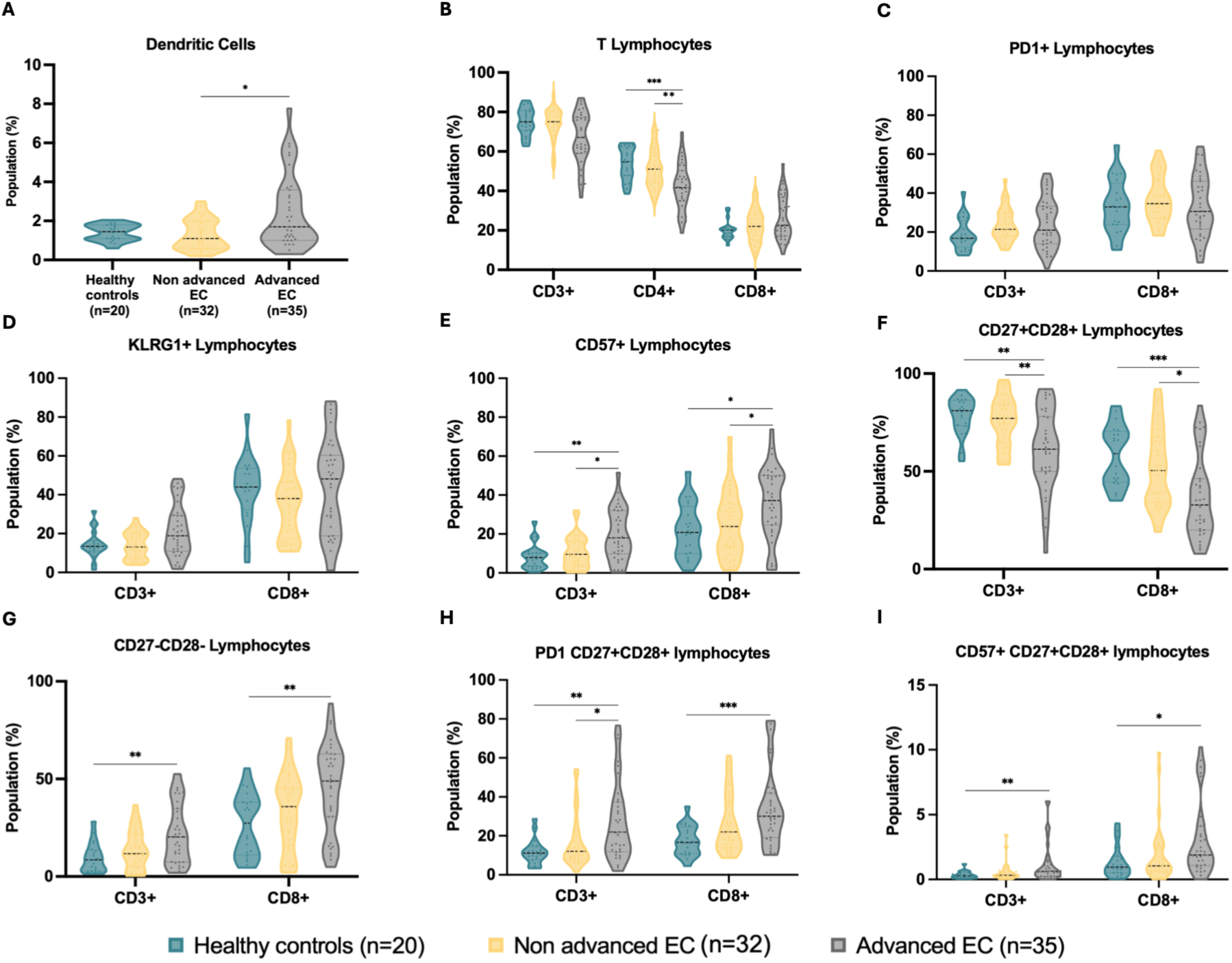
Immune cell profiles in EC patients. Levels of different immune cell populations in healthy controls, localized and advanced endometrial tumors: **A.** Dendritic cells, **B.** CD3^+^/CD4^+^/CD8^+^ cells, **C.** PD1^+^, **D.** KLRG1^+^, **E.** CD57^+^ T cells, **F**. CD27^+^CD28^+^, **G.** CD27^-^CD28^-^, **H-I.** PD1^+^/CD57^+^ within CD27^+^CD28^+^ and CD27^-^CD28^-^ compartments of CD3^+^ and CD8^+^ subsets. *p < 0.05, **p < 0.01, ***p < 0.001, Kruskal-Wallis test with Dunn’s post-hoc analysis.

Within the lymphoid lineage, both CD4^+^ and CD8^+^ T lymphocytes were characterized among the global population of CD3^+^ cells. CD4^+^ cells were significantly reduced in advanced-stage tumors compared to localized tumors (post hoc Dunn’s adjusted *p* = 0.008) and controls (post hoc Dunn’s adjusted *p* = 7.0×10^-4^) (Figure 1.B). By contrast, global CD8^+^ lymphocyte proportions did not differ significantly between groups.

Cells expressing PD1, KLRG1 and CD57 were also studied among the lymphoid lineage to better understand tumor immunity and cellular senescence. No significant differences were observed in PD1^+^ T cells between groups after adjustment for multiple testing (Figure 1.C). Similarly, the KLRG1^+^ population did not retain statistical significance following correction (Figure 1.D). In contrast, CD8^+^CD57^+^ cells were significantly increased in patients with advanced EC compared to localized EC and healthy individuals (post hoc Dunn’s adjusted *p* = 0.026 and 0.013, respectively) (Figure 1.E), supporting the hypothesis of a potential role of senescent or terminally differentiated T cells in disease progression.

To further delineate the immune landscape, we characterized lymphocyte subpopulations based on CD27 and CD28 expression, two key co-stimulatory molecules involved in immune modulation. CD27^+^CD28^+^ lymphocytes were decreased in advanced disease, particularly within the cytotoxic lymphocyte subset (global *p* = 0.004) (Figure 1.F). Conversely, senescent-like CD27^-^CD28^-^ lymphocytes were significantly increased in advanced disease (CD8^+^ global *p* = 0.0036) (Figure 1.G). Together, these findings suggest a shift in cytotoxic T-cell composition in advanced disease.

Cells expressing PD1, KLRG1 and CD57 were also studied within CD27^+^CD28^+^ and CD27^-^CD28^-^ lymphocyte populations to better understand tumor immunity and cellular senescence. We observed a highly significant increase in PD-1 expression, consistent with an exhaustion-associated phenotype, in CD3^+^ and CD8^+^ CD27^+^CD28^+^ lymphocytes in all EC patients (global *p* = 0.003 and 8.0×10^-4^, respectively) (Figure 1.H). Furthermore, we also noted a significant increase in CD57 presence in patients with advanced tumors (global *p* = 0.007 and 0.02, respectively) (Figure 1.I). Further comparisons conducted within these groups are depicted in Supplementary Figure 3.

Overall, the comparison of immune populations suggests that the most pronounced changes occur in advanced disease, whereas localized tumors largely resemble the control group. These patterns are consistent with a disease-associated shift in circulating immunity.

### Association between Peripheral Immune Population Levels and Clinicopathologic characteristics

To explore the clinical relevance of immune cell profiles identified, we analyzed associations between immune populations and key clinicopathologic variables, including EC type, grade, myometrial infiltration, MSI status and TP53 status. Among the clinicopathologic variables analyzed, tumor grade showed a significant association with MDSC levels. Specifically, G2 tumors exhibited a higher proportion of MDSC-high cases compared with both G1 and G3 tumors (*p* = 0.05) (Figure 2.A, Supplementary Table 2). EC subtype was significantly associated with CD8^+^CD27^+^CD28^+^ levels (*p* = 0.018) and MDSCs (*p* = 0.039) (Figures 2.B-C, Supplementary Table 3).

Myometrial infiltration was also associated with MDSC levels, with tumors showing >50% infiltration displaying a higher proportion of MDSC-high cases than tumors with ≤50% infiltration (p = 0.019) (Figure 2.D, Supplementary Table 4). Although this association does not establish causality, it is consistent with the possibility that deeper local invasion may be accompanied by a more immunosuppressive peripheral profile.

**Figure 2.**
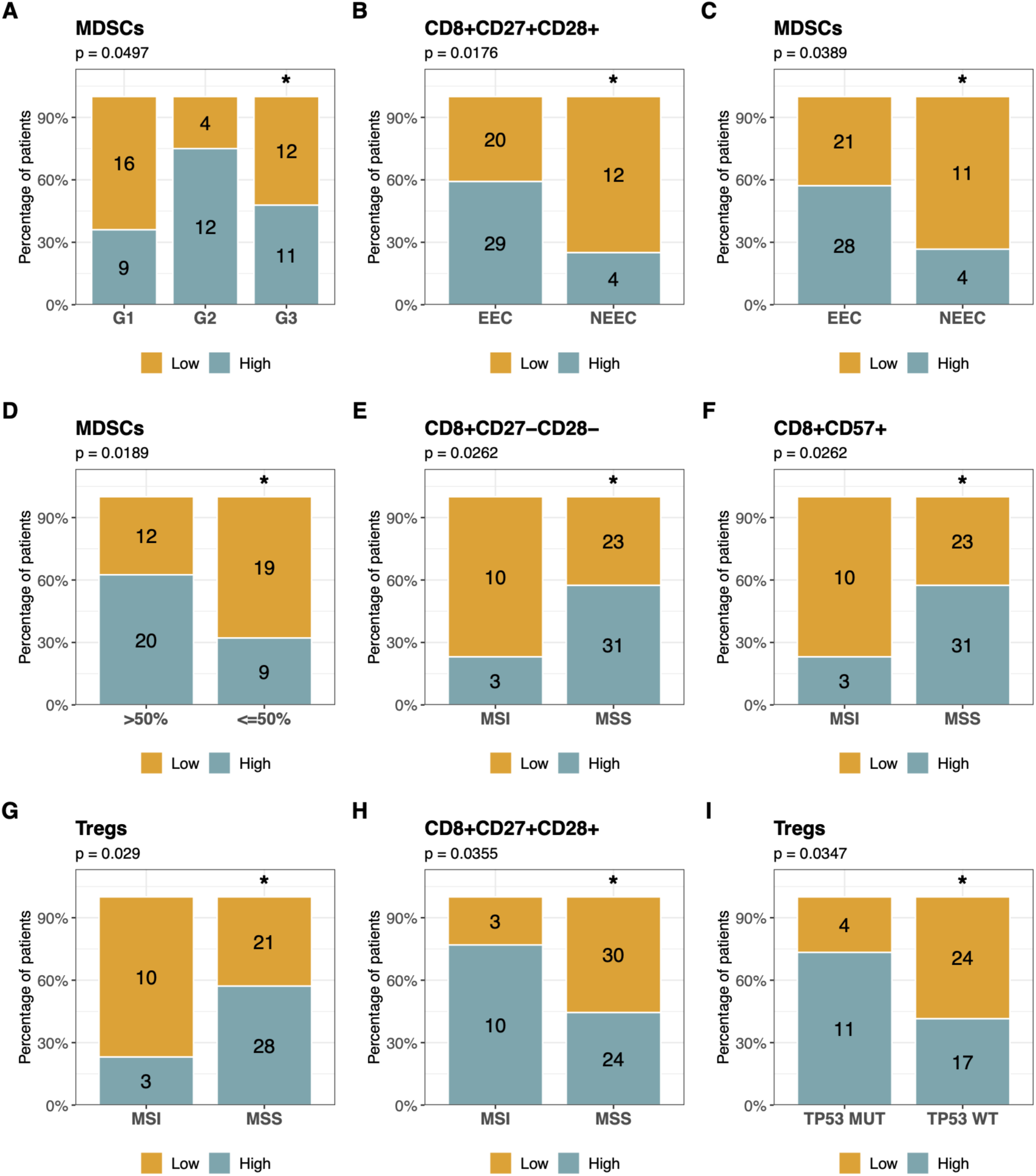
Clinicopathological associations of immune subsets. Stacked bar plots show the proportion of low and high immune cell levels within each clinicopathological category. Numbers inside the bars indicate case counts, and significance is denoted as *p<0.05, **p<0.01 and ***p<0.001.

Regarding molecular features, MSS tumors were associated with high CD8^+^CD27^-^CD28^-^ (*p* = 0.026), CD8^+^CD57^+^ (*p* = 0.026) and Tregs (*p* = 0.029), as well as lower levels of CD8^+^CD27^+^CD28^+^ (*p* = 0.036) (Figures 2.E-H, Supplementary Table 5). Together, these findings suggest that MSS tumors may display a distinct peripheral immune phenotype involving both regulatory and differentiated T-cell compartments. Finally, *TP53* status was associated with Tregs (*p* = 0.035), with higher Treg levels in mutated tumors (Figure 2.I, Supplementary Table 6). This finding may reflect a more immunosuppressive peripheral environment in tumors harboring *TP53*alterations.

Overall, these results suggest that distinct clinicopathologic EC subgroups may be associated with subtle but biologically meaningful differences in the peripheral immune compartment.

### Association Between Baseline Peripheral Blood Immune Cell Levels and Disease Progression in Advanced EC Patients

To assess the potential utility of our immunophenotyping strategy in predicting EC patient outcomes, we analyzed the association between baseline immune cell populations and disease evolution in the advanced-disease cohort. To accomplish this, baseline immune profiles were analyzed in 31 patients with advanced EC who had at least 6 months of follow-up from treatment onset. Patients without progressing disease after the treatment onset were defined as responders (R, n=16) while those progressing before 6 months were classified as early PD (n=7) and after 6 months as late PD (n=8). Of these, 21 received first-line chemotherapy, 7 received first-line hormone therapy, and 3 received first-line immunotherapy. Treatments of these patients are shown in Figure 3.A.

**Figure 3.**
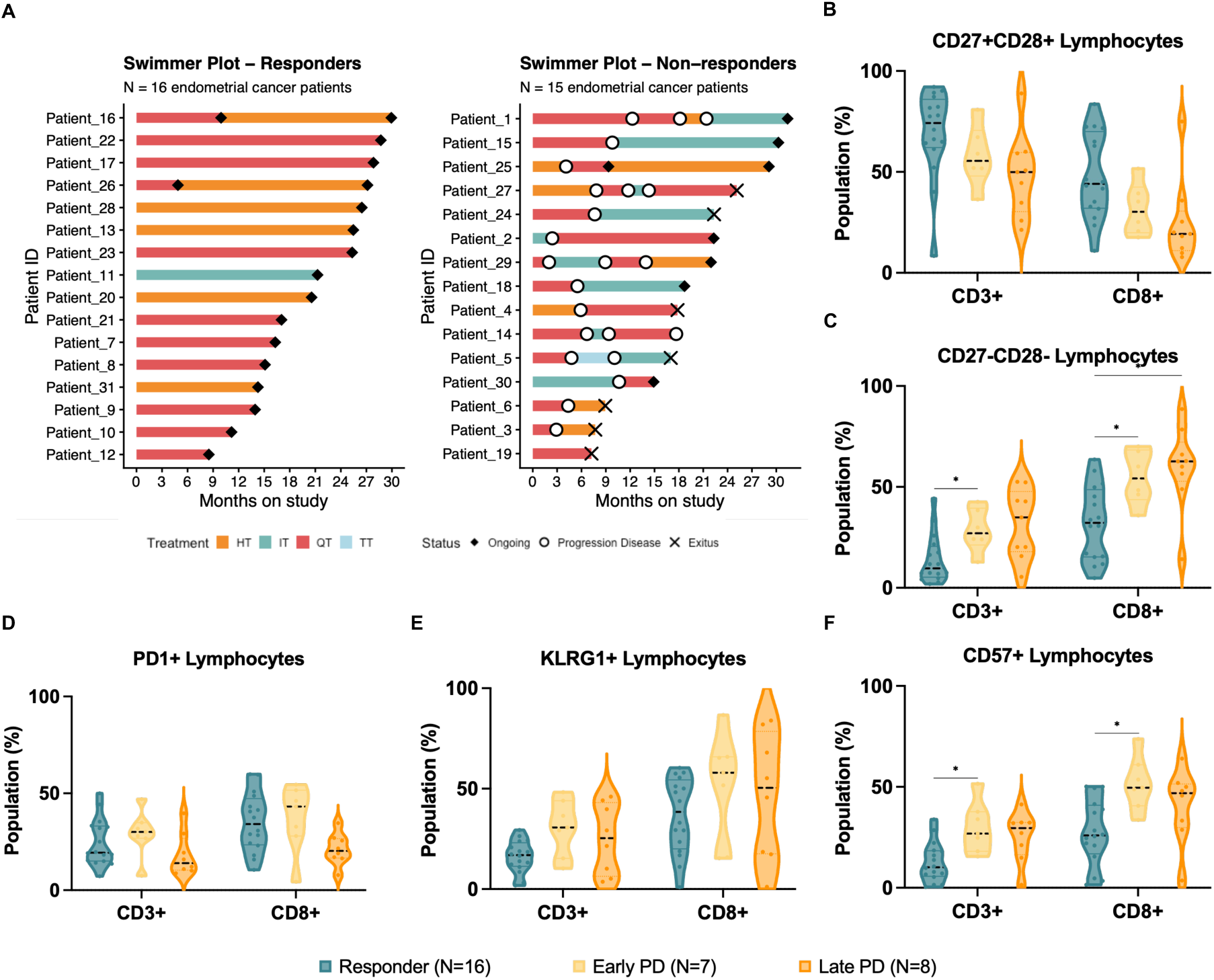
Integrated Analysis of Immune Cell Subtypes and Treatment Outcomes in EC Patients. **A.** Swimmer plots depicting follow-up data from our cohort. HT: Hormone therapy; IT: Immunotherapy; QT: Chemotherapy; TT: Targeted therapy. **B.** Comparison of CD27^+^CD28^+^ lymphocytes between R, early PD and late PD patients. **C.** CD27^-^CD28^-^ lymphocytes. **D-F.** PD1, KLRG1 and CD57 levels.

Both groups of PD patients showed lower levels of CD27^+^CD28^+^ lymphocytes, although it did not reach statistical significance (Figure 3.B). Notably, early and late progressors also showed higher levels of CD27^-^CD28^-^ lymphocytes (R vs. Early PD for CD3^+^: adjusted *p* = 0.026; R vs. Early PD for CD8*^+^*: adjusted *p* = 0.033; R vs. Late PD for CD8^+^: adjusted *p* = 0.016) (Figure 3.C). No statistically significant differences were found in PD1 and KLRG1 levels (Figures 3.D-E). However, we observed a significant increase in CD3^+^ and CD8^+^CD57^+^ lymphocytes in early PD (adjusted *p* = 0.016 and 0.013, respectively) (Figure 3.F). These findings suggest an increase in more differentiated lymphocyte subsets in patients with early progression. Further comparisons are shown in Supplementary Figure 4.

The impact of the distinct immune cell populations on PFS was also examined. Kaplan-Meier analysis revealed that patients with high baseline levels of total CD8^+^ lymphocytes had significantly worse PFS (HR 3.62, 95% CI 1.15–11.43; log-rank *p* = 0.019) (Figure 4.A). In contrast, patients with higher levels of CD27^+^CD28^+^ lymphocytes showed improved PFS (Figure 4.B). Higher proportions of both CD27^-^CD28^-^ lymphocytes (HR 5.1, 95% CI 1.43– 18.17; log-rank *p* = 0.005) (Figure 4.C) and CD8^+^CD57^+^ lymphocytes (HR 3.73, 95% CI 1.18–11.78; log-rank *p* = 0.016) (Figure 4.D) were likewise associated with poorer PFS.

**Figure 4.**
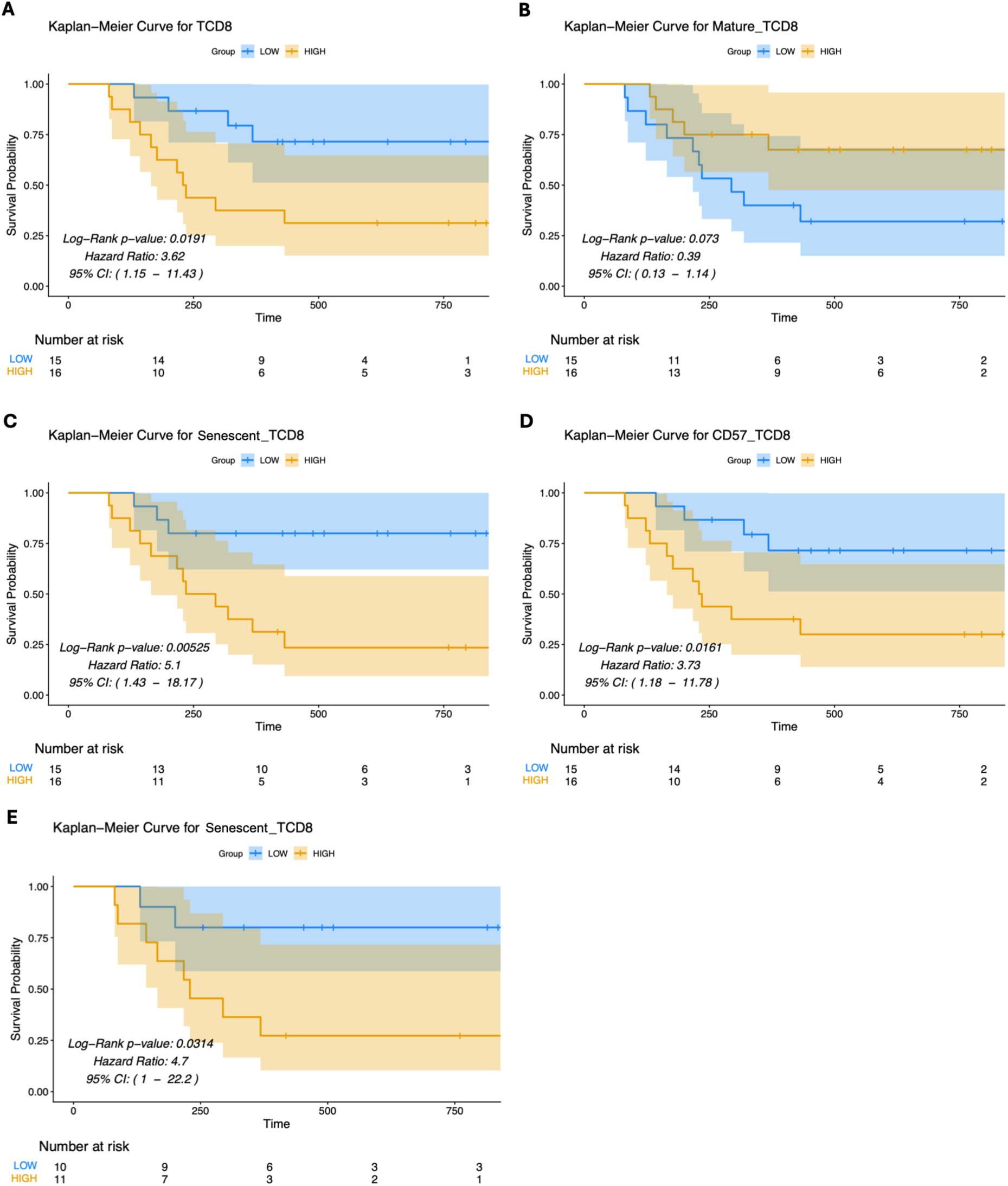
Prognostic Impact of Immune Cell Subtypes on Treatment Outcomes in EC Patients. **A.** Kaplan-Meier plot of PFS among patients with high levels of CD8^+^ lymphocytes. **B.** CD8^+^CD27^+^CD28^+^ lymphocytes. **C.** CD8^+^CD27^-^CD28^-^ lymphocytes and **D.** CD57^+^ lymphocytes. **E.** CD8^+^CD27^-^CD28^-^ lymphocytes in patients receiving chemotherapy as 1^st^ line treatment.

Exploratory univariate analysis revealed that non-responders more frequently displayed elevated CD8^+^, CD8^+^CD27^-^CD28^-^, and CD8^+^CD57^+^ lymphocyte levels, and decreased mature CD8^+^ (Supplementary Table 7). Results for the remaining immune populations are shown in Supplementary Figure 5. In the univariate Cox analysis, higher baseline proportions of total CD8^+^ cells (HR 1.057, 95% CI 1.016-1.099, *p* = 0.006), CD8^+^CD27^-^ CD28^-^ cells (HR 1.043, 95% CI 1.015-1.072, *p* = 0.003), and CD8^+^CD57^+^ cells (HR 1.063, 95% CI 1.020-1.108, *p* = 0.004) were significantly associated with worse PFS. The combined low mature/high senescent CD8^+^ phenotype also showed a significant association with poorer outcome (HR 1.014, 95% CI 1.003-1.025, *p* = 0.015). In the exploratory multivariable Cox model adjusted for age, histology, grade, and myometrial infiltration, CD8^+^ (HR 1.134, 95% CI 1.049-1.227, *p* = 0.002), CD8^+^CD27^-^CD28^-^ (HR 1.039, 95% CI 1.009-1.071, *p* = 0.011), and CD8^+^CD57^+^ (HR 1.069, 95% CI 1.021-1.119, *p* = 0.004) were independently associated with shorter PFS. (Supplementary Tables 8-9).

Survival analyses restricted to the 21 patients who received chemotherapy for advanced disease were also performed to account for potential effects of treatment regimen on PFS. For CD8^+^CD27^-^CD28^-^ lymphocytes, increased levels were again associated with worse PFS (HR 4.7, 95% CI 1-22.2; log-rank *p* = 0.032). Kaplan-Meier analyses of the remaining cells are shown in Supplementary Figure 6.

Spearman correlation analysis in the advanced cohort supported a coordinated remodeling of the CD8 compartment, with the expected inverse relationship between mature and late-differentiated CD8 subsets and positive associations between senescent markers and terminally differentiated phenotypes. This pattern is consistent with the clinical associations observed for response and PFS (Supplementary Figure 7).

Collectively, these findings suggest that a baseline systemic immune profile characterized by senescent T cells may be associated with poor treatment response and worse PFS, consistent with more aggressive tumor behavior.

### Longitudinal Evolution of Circulating Immune Cell Populations and Association with Disease Progression

To elucidate the dynamics of immune responses during therapy, we performed a longitudinal analysis of peripheral blood immune cell subpopulations in 28 patients from our study with longitudinal samples. CD8^+^ cells showed a separation between groups over the treatment course. While no significant differences were observed at baseline or 3 months, patients with high CD8^+^ proportions at 6 months were more likely to develop progressive disease (Fisher’s exact test, *p* = 0.015; OR = 18.80, 95% CI: 1.31-1242.44) (Figure 5.A). Mature lymphocytes showed the opposite pattern since non-progressive patients maintained higher proportions throughout the follow-up (Supplementary Figure 8.A). At baseline, low mature lymphocyte levels tended to associate with progressive disease (*p* = 0.057; OR = 0.11, 95% CI: 0.01-1.27), although this did not reach statistical significance.

**Figure 5.**
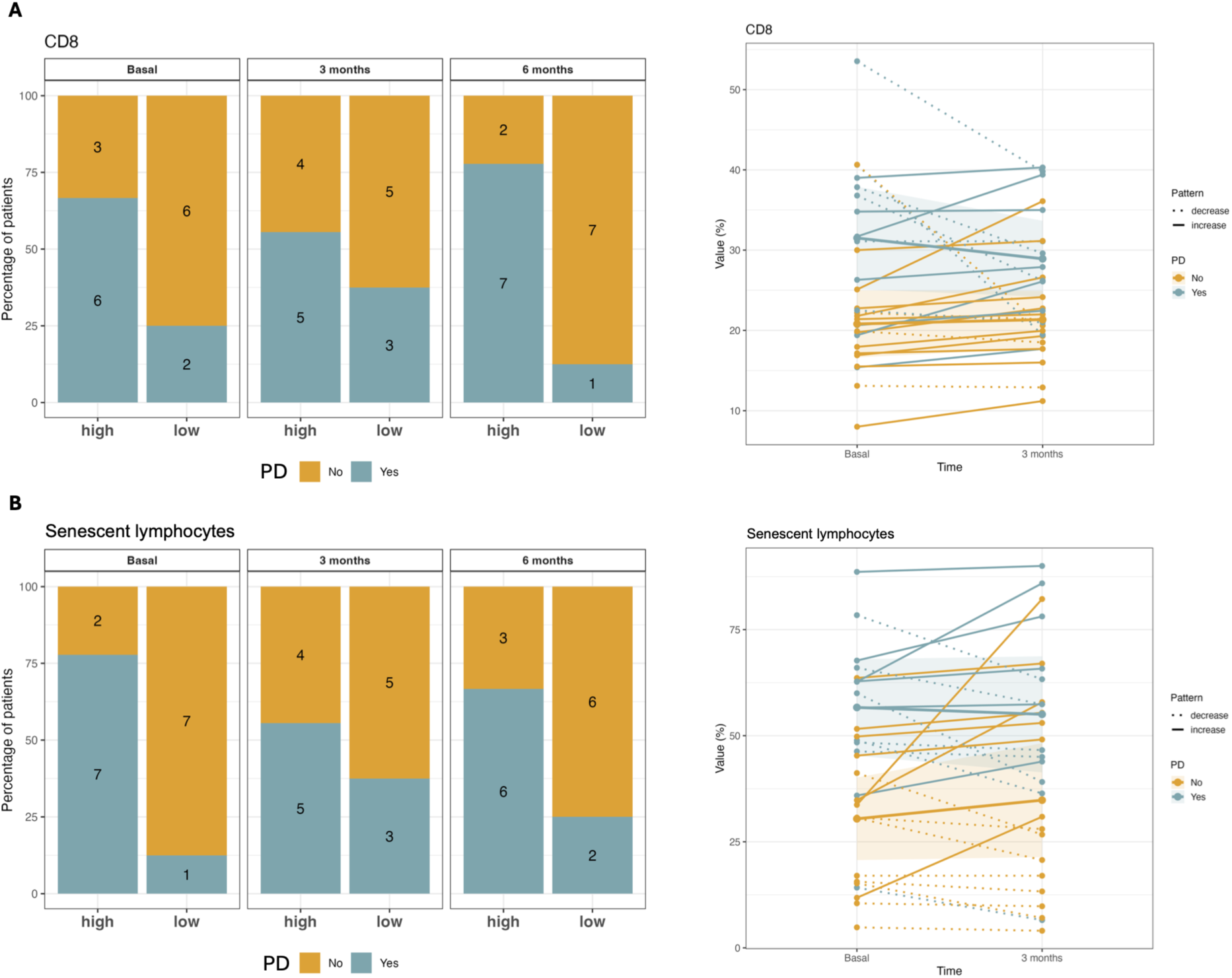
Longitudinal analysis of peripheral blood immune cell subpopulations with advanced EC. **A.** Total CD8^+^ lymphocytes. **B.** Senescent-like lymphocytes. Barplots (left panels) show the percentage of patients with high and low values at baseline, 3 and 6 months. Paired-line plots (right panels) illustrate individual trajectories for each cell subset, stratified by disease progression.

In addition, senescent-like lymphocytes showed the strongest association with disease evolution. High baseline and follow up levels were associated with disease progression (*p* = 0.015; OR = 18.80, 95% CI: 1.31-1242.44; Figure 5.B), suggesting a stable difference in immune profiles over time. Finally, CD8^+^CD57^+^ cells showed a consistent but non-significant trend towards higher proportions in PD patients at all timepoints (*p* = 0.153; OR = 5.33, 95% CI: 0.52-87.73; Supplementary Figure 8.B), with a remarkably stable distribution pattern throughout the follow-up. These findings are consistent with the role of terminally differentiated CD57^+^ cytotoxic T cells in immune senescence and tumor immune evasion.

## DISCUSSION

Immune cells play a crucial role in halting the progression of solid tumors by identifying and eliminating tumor cells [12]. Nonetheless, comprehensive analyses of peripheral circulating immune cells in EC patients and their potential implications for tumor prognosis are still lacking. The results obtained in the present study provide exploratory insight into the complex interplay between immune cell populations and clinical outcomes in advanced EC. Our findings reveal notable differences across disease stages, suggesting that the immune microenvironment may be associated with disease progression and treatment efficacy. These insights emphasize the necessity of incorporating immune factors into the comprehensive assessment of EC to better inform therapeutic strategies and improve patient outcomes.

In our study, the comparison of the immune composition between healthy controls and patients with localized and advanced EC revealed significant differences between the advanced disease and the other groups, suggesting pronounced regulation in advanced stages. These included higher dendritic cell levels, lower CD4^+^ T cell proportions, and an increased frequency of CD3^+^CD57^+^ cells. In addition, advanced-stage EC patients exhibited greater signs of immunosenescence, reflected by elevated CD57^+^ T cells and CD8^+^CD27^-^CD28^-^ populations. The increased proportion of dendritic cells suggests a potential role for these cells in tumor progression, consistent with previous studies in other malignancies [16, 17]. Although dendritic cells can activate strong antitumor T cell responses, tumor-derived factors may exploit them to facilitate cancer progression [18]. Of note, high levels of this population have been associated with a poor prognosis in other types of tumors, such as hepatocellular carcinoma [16]. In particular, plasmacytoid dendritic cells may contribute to an immunosuppressive tumor microenvironment by inhibiting cytotoxic T cell activity and recruiting regulatory T cells. The concurrent decrease in CD4^+^ lymphocytes in advanced disease further highlights the dynamic reshaping of the peripheral immune landscape [19, 20], in line with reports associating low CD4^+^ levels at diagnosis with poor therapy response in other tumors [21].

EC cells employ various strategies to circumvent immune surveillance, contributing to tumor progression and resistance. In our study, CD57 emerged as the most consistent marker of a senescent-like phenotype, particularly in advanced disease and in specific CD8^+^ subsets. By contrast, PD-1 and KLRG1 showed more limited or subset-restricted associations, suggesting that their relevance may depend on the phenotypic context rather than reflecting a generalized increase across all T-cell compartments. Endometrial tumors often exploit the interaction between PD-1/PD-L1 axis to promote T cell dysfunction, which represents the principal immunotherapy target in this malignancy [6, 22]. The increased levels of CD57 in selected T-cell compartments support the presence of senescent-like subsets, while the PD-1 and KLRG1 findings should be interpreted as restricted to specific phenotypic contexts rather than as global shifts across the entire T-cell pool [23, 24]. CD57 identifies senescent-like cells with reduced proliferative capacity, though its impact differs between T cells and NK cells [25]. Moreover, while CD57^+^ NK cells have been associated with less severe outcomes in cancer patients, the presence of CD8^+^CD57^+^ cells has been linked to poor OS, underlining their role as negative prognostic markers in advanced malignancies [26, 27].

Moreover, the observed associations between tumor molecular characteristics and peripheral immune cell subsets support the concept that tumor-intrinsic alterations contribute to shaping systemic immune responses. The immune profile associated with MSS tumors is consistent with the lower immunogenicity typically observed in MSS tumors and could contribute to the limited efficacy of immunotherapy in this molecular subtype through the establishment of an immunosuppressive systemic environment. These findings are in agreement with current models of T-cell dysfunction in cancer and the central role of regulatory T cells in suppressing antitumor immunity [28, 29].

The association between *TP53* alterations and increased circulating regulatory T cells further supports the emerging role of TP53 as a regulator of tumor-immune interactions beyond its canonical functions in cell cycle control and genomic stability. Increasing evidence indicates that *TP53* mutations promote immune evasion through multiple mechanisms, including modulation of inflammatory signaling, impaired antigen presentation, and the establishment of an immunosuppressive microenvironment enriched in suppressive immune cell populations [30]. Although the mechanisms underlying the present observations require further investigation, our findings are consistent with these emerging immunomodulatory functions of *TP53*.

Furthermore, the longitudinal analysis reinforced these baseline findings. High baseline proportions of senescent-like CD27^-^CD28^-^ lymphocytes were associated with disease progression, and this separation was maintained throughout the entire follow-up, indicating a stable immune dysfunction consistent with T cell senescence as a mechanism of immune evasion in solid tumors [31]. Notably, CD8^+^ T cell proportions were also elevated in progressive patients at 6 months, which may reflect an accumulation of dysfunctional cytotoxic T cells unable to mount an effective anti-tumour response, a phenomenon described in the context of checkpoint inhibitor resistance [32]. In contrast, non-progressive patients maintained higher mature lymphocyte proportions throughout follow-up, supporting the role of a functionally competent lymphocyte compartment in sustained treatment benefit.

Importantly, the association between baseline immune cell levels and clinical outcomes underscores its potential prognostic value in guiding therapeutic decisions. Notably, CD8^+^CD27^-^CD28^-^ was the marker whose association with outcome was most consistent across the different analytical approaches used in this study, including the categorical relapse association, the univariate Cox model, and the multivariable Cox model, supporting it as a robust candidate biomarker identified here. Patients with higher levels of senescent lymphocytes exhibited poorer PFS rates, emphasizing the need for targeted interventions to overcome immune dysfunction [33]. Conversely, patients with higher levels of CD27^+^CD28^+^ lymphocytes demonstrated improved outcomes, suggesting a protective role for these mature functional immune cells [34, 35].

Nevertheless, our findings should be interpreted in light of several limitations. The relatively small sample size may limit generalizability, and the heterogeneity of treatment regimens may complicate direct comparisons of immune responses. In addition, categorical analyses were based on median dichotomization, an approach that may be sensitive to sample size and cohort-specific distributions and may not reflect biologically optimal thresholds. Finally, the longitudinal and multivariable analyses were exploratory, and the limited number of events increases the risk of overfitting and unstable effect estimates.

In conclusion, this study provides a comprehensive characterization of the peripheral immune landscape across the clinical spectrum of EC and demonstrates that advanced disease is accompanied by distinct systemic immune remodeling. The identification of circulating immune populations associated with disease stage, progression, and clinical outcome highlights the potential of peripheral immune profiling to capture biologically and clinically relevant features of the host immune response. Beyond improving our understanding of systemic immunity in endometrial cancer, these findings provide a framework for the development of minimally invasive immune biomarkers for patient stratification and disease monitoring. Although confirmation in larger prospective cohorts is warranted, this work establishes a valuable resource for future translational studies investigating the role of systemic immunity in EC.

## Conflicts of Interest

The authors declare that they have no conflict of interest.

## Funding

This work was supported by grants and support from the Instituto de Salud Carlos III (ISCIII) (PI21/00990), FEDER, CIBERONC (CB16/12/00328) and GAIN Proyectos de Excelencia (IN607D2021/05) to L.M-R. In addition, L.M-R. is supported by a contract “Miguel Servet” from ISCIII (CP20/00119). Additionally, R.P-P. is supported by a predoctoral fellowship from ISCIII (FI22/00207).

## Author Contributions

Conceptualization: R.P-P, L.M-R, J.V-R; Methodology: R.P-P, L.M-R, J.V-R; Investigation (patient recruitment, clinical data collection, sample collection, and processing): A.V, E.A, V.S, A.A, C.R, A.C, R.M, E.D, I.P, S.B-P, L.V-T, I.F-P, E.M-M, S.C, C.C, A.H, R.L-L, J.C, R.P-P, J.V-R; Resources: A.V, E.A, V.S, A.A, C.R, A.C, R.M, E.D, I.P, S.B-P, L.V-T, I.F-P, E.M-M, S.C, C.C, A.H, R.L-L J.C, E.M-M; Data curation: R.P-P, J.V-R; Formal analysis: R.P-P, J.V-R; Writing - original draft: R.P-P, L.M-R; Writing - reviewing & editing: J. V-R, G.M-B, S.C; Funding acquisition: L.M-R.

## Ethics approval

This study received approval from the Galician Autonomic Ethics Committee (Comité Autonómico de Ética de la Investigación de Galicia, Xunta de Galicia, Spain; Ref: 2022/029), which served as the central ethics committee. The study was conducted in accordance with Good Clinical Practice guidelines and the Declaration of Helsinki. Prior to participation, all patients provided written informed consent.

## Data Availability Statement

Data is accessible upon reasonable request from the corresponding author.

## Acknowledgments

This study would not have been possible without the kind collaboration of all the patients. We also thank Inma Hernández, nurse at the Hospital General Universitario de Valencia (CHGUV), for her exceptional efforts and invaluable support in collecting the ENDOPERC study samples.

## Conflicts of Interest

The authors declare that they have no conflict of interest.

## Notes

### Competing Interest Statement

The authors have declared no competing interest.

### Summary of Updates

Upload with the Figure 3 revised.

