## Supplementary Files for "Circulating Immune Profiling Reveals Immune Signatures Associated with Disease Stage and Outcome in Endometrial Cancer"

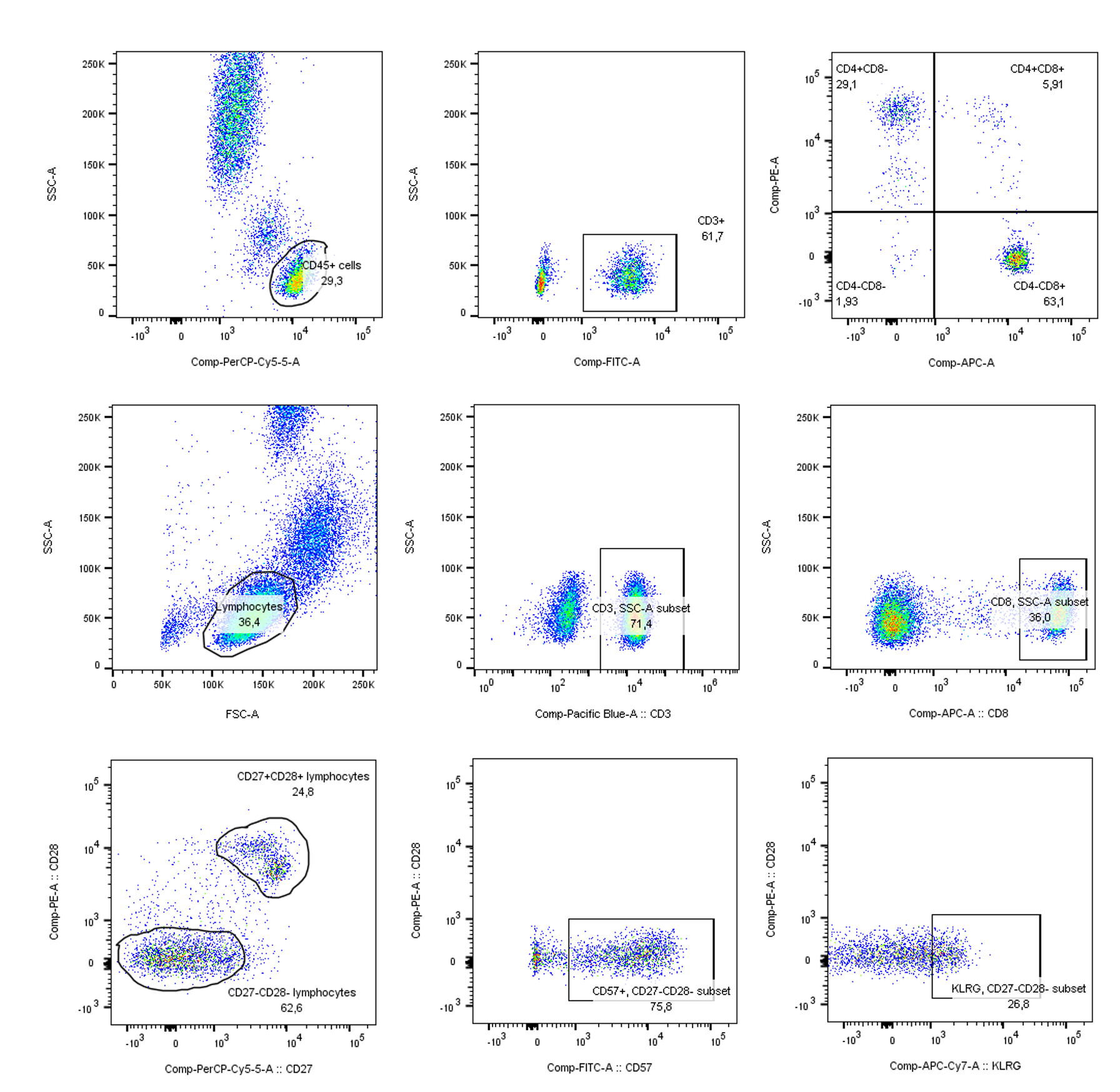

**Supplementary Figure 1. Immune cell analysis in peripheral blood of patients with EC.** Lymphocytes were identified and gated based on FSC-A vs. SSC-A, followed by selection of CD3^+^, CD4^+^ and CD8^+^ T cells. Further characterization of memory and senescent subsets was performed using CD27^+^, CD28^+^, CD57^+^ and KLRG1^+^ expression.

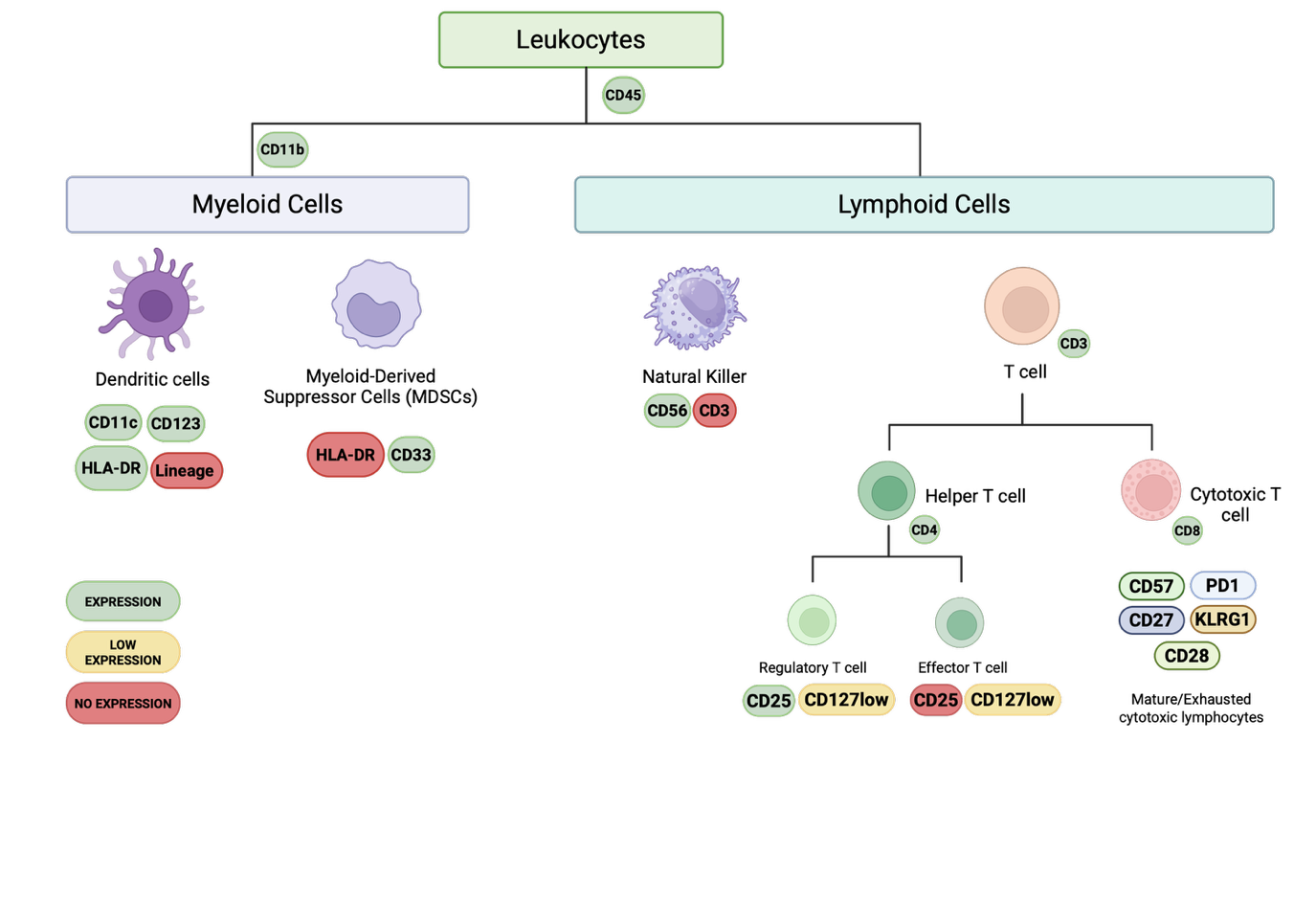
**Supplementary Figure 2.** Illustration of immune cell populations studied and the specific antibodies employed for their identification. Font: Biorender.

**Supplementary Table 1. List of Antibodies Used for Flow Cytometry Analysis.** Details of the antibodies used in the study, including marker, the specific cell populations tagged, fluorochrome label, host species, target species, brand and the catalog number.

| **Marker** | **Targeted Cell Type** | **Label** | **Host** | **Target** | **Brand** | **Catalog No.** | **Volume used for analysis** |
| --- | --- | --- | --- | --- | --- | --- | --- |
| **CD3** | T lymphocytes | FITC | Mouse | Human | BD | 340542 | 3 µL |
| **CD14** | Monocytes | FITC | Mouse | Human | BD | 345784 | 3 µL |
| **CD19** | B lymphocytes | FITC | Mouse | Human | BD | 340864 | 3 µL |
| **CD56** | NK cells | FITC | Mouse | Human | BD | 345811 | 3 µL |
| **HLA-DR** | APC | PerCP | Mouse | Human | BD | 347364 | 10 µL |
| **CD11c** | Dendritic cells | APC | Mouse | Human | BD | 333144 | 5 µL |
| **CD123** | Plasmacytoid DC | PE | Mouse | Human | Biolegend | 306006 | 5 µL |
| **CD11b** | Monocytes, granulocytes | PE | Mouse | Human | BD | 347557 | 10 µL |
| **CD33** | Myeloid cells | APC | Mouse | Human | BD | 551378 | 10 |
| **CD127** | Memory T cells | APC | Mouse | Human | Biolegend | 351316 | 5 µL |
| **CD25** | Regulatory T cells | PE | Mouse | Human | Biolegend | 302606 | 5 µL |
| **CD4** | Helper T cells | FITC | Mouse | Human | Biolegend | 357406 | 5 µL |
| **CD45** | Hematopoietic cells | PerCP | Mouse | Human | BD | 342410 | 10 µL |
| **CD3** | T lymphocytes | FITC | Mouse | Human | BD | 342410 | 10 µL |
| **CD4** | Helper T cells | APC | Mouse | Human | BD | 345771 | 5 µL |
| **CD8** | Cytotoxic T cells | PE | Mouse | Human | BD | 342410 | 10 µL |
| **CD3** | T lymphocytes | PerCP | Mouse | Human | BD | 347344 | 10 µL |
| **CD19** | B lymphocytes | APC | Mouse | Human | BD | 340722 | 5 µL |
| **CD56** | NK cells | PE | Mouse | Human | BD | 345812 | 10 µL |
| **CD3** | T lymphocytes | V450 | Mouse | Human | BD | 560365 | 5 µL |
| **CD57** | Senescent T cells | FITC | Mouse | Human | BD | 347393 | 10 µL |
| **CD27** | Memory T cells | PerCP | Mouse | Human | BD | 560612 | 10 µL |
| **CD28** | Activated T cells | PE | Mouse | Human | BD | 561793 | 10 µL |
| **CD8** | Cytotoxic T cells | APC | Mouse | Human | BD | 561952 | 5 µL |
| **CD279** | Regulatory, exhausted T cells | PE-Cy7 | Mouse | Human | Biolegend | 561272 | 5 µL |
| **KLRG1** | Senescent T cells | APC-Cy7 | Syrian Hamster | Mouse/Human | Biolegend | 138426 | 5 µL |

**A**

**B**

**C**

**D**

**E**

**F**

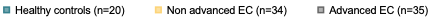

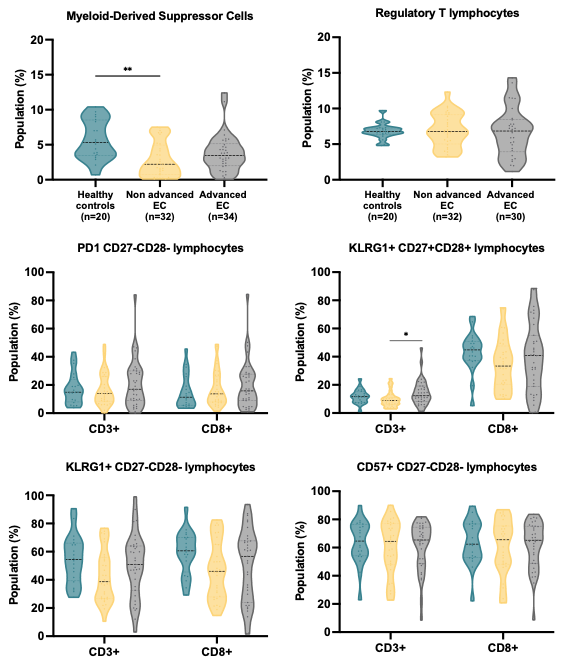

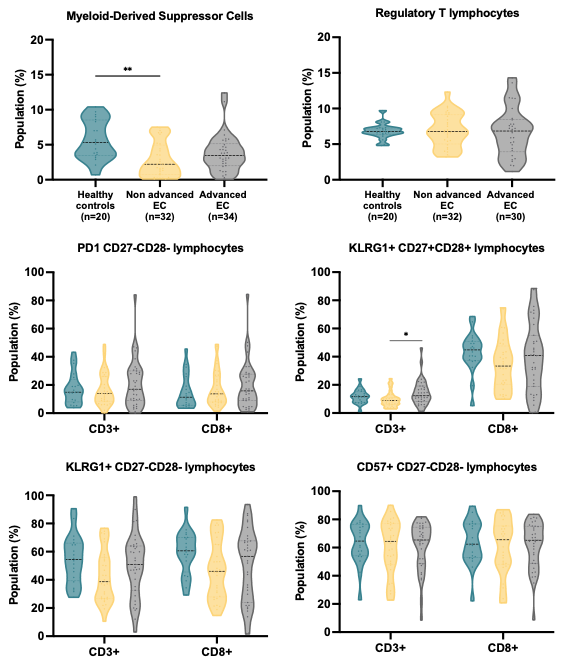

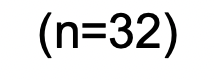

**Supplementary Figure 3. Immune cell profiles in healthy controls and EC patients. A.** MDSCs levels across healthy controls, patients with localized and advanced disease. **B.** Tregs levels. **C.** PD1 expression in CD27^-^CD28^-^ lymphocytes. **D.** KLRG1 expression in CD27^+^CD28^+^ lymphocytes. **E.** KLRG1 expression in CD27^-^CD28^-^ lymphocytes. **F.** CD57 expression in CD27^-^CD28^-^ lymphocytes. Data are presented as mean ± standard deviation. **p* < 0.05, ***p* < 0.01, ****p* < 0.001, Kruskal-Wallis test with Dunn's post-hoc analysis.

**Supplementary Table 2.** **Distribution of the peripheral immune cell profiles based on the tumor grade.**

| **Grade** |  | **G1** N = 25*^1^* | **G2** N = 16*^1^* | **G3** N = 24*^1^* | **p-value***^2^* | **q-value***^3^* |
| --- | --- | --- | --- | --- | --- | --- |
| **Dendritic_cells** | Low | 11 (44%) | 9 (56%) | 11 (46%) | 0.7 | 0.774 |
|  | High | 14 (56%) | 7 (44%) | 13 (54%) |  |  |
| **MDSCs** | Low | 16 (64%) | 4 (25%) | 12 (52%) | **0.050** | 0.303 |
|  | High | 9 (36%) | 12 (75%) | 11 (48%) |  |  |
| **Tregs** | Low | 14 (56%) | 8 (53%) | 9 (45%) | 0.8 | 0.774 |
|  | High | 11 (44%) | 7 (47%) | 11 (55%) |  |  |
| **CD8^+^** | Low | 11 (44%) | 11 (69%) | 10 (42%) | 0.2 | 0.489 |
|  | High | 14 (56%) | 5 (31%) | 14 (58%) |  |  |
| **NK** | Low | 16 (70%) | 7 (44%) | 9 (39%) | 0.091 | 0.303 |
|  | High | 7 (30%) | 9 (56%) | 14 (61%) |  |  |
| **CD8^+^CD27^+^CD28^+^** | Low | 10 (40%) | 7 (44%) | 14 (58%) | 0.4 | 0.715 |
|  | High | 15 (60%) | 9 (56%) | 10 (42%) |  |  |
| **CD8^+^CD27^-^CD28^-^** | Low | 14 (56%) | 8 (50%) | 11 (46%) | 0.8 | 0.774 |
|  | High | 11 (44%) | 8 (50%) | 13 (54%) |  |  |
| **CD8^+^PD1^+^** | Low | 12 (48%) | 6 (38%) | 14 (58%) | 0.4 | 0.715 |
|  | High | 13 (52%) | 10 (63%) | 10 (42%) |  |  |
| **CD8^+^KLRG1^+^** | Low | 14 (56%) | 7 (44%) | 11 (46%) | 0.7 | 0.774 |
|  | High | 11 (44%) | 9 (56%) | 13 (54%) |  |  |
| **CD8^+^CD57^+^** | Low | 17 (68%) | 6 (38%) | 10 (42%) | 0.087 | 0.303 |
|  | High | 8 (32%) | 10 (63%) | 14 (58%) |  |  |
| *^1^* n (%) |  | | | | |  |
| *^2^* Pearson’s Chi-squared test  *^3 Benjamini & Hochberg correction for multiple testing^* |  | | | | |  |
| *^3^* Benjamini & Hochberg correction for multiple testing |  | | | | |  |

**Supplementary Table 3. Distribution of the peripheral immune cell profiles between EC histologic subtypes.**

| **Type** |  | | **EEC** N = 49*^1^* | **NEEC** N = 16*^1^* | **p-value***^2^* | **q-value***^3^* |
| --- | --- | --- | --- | --- | --- | --- |
| **Dendritic_cells** | Low | | 22 (45%) | 9 (56%) | 0.4 | 0.614 |
|  | High | | 27 (55%) | 7 (44%) |  |  |
| **MDSCs** | Low | | 21 (43%) | 11 (73%) | **0.039** | 0.194 |
|  | High | | 28 (57%) | 4 (27%) |  |  |
| **Tregs** | Low | | 24 (53%) | 7 (47%) | 0.7 | 0.818 |
|  | High | | 21 (47%) | 8 (53%) |  |  |
| **CD8^+^** | Low | | 23 (47%) | 8 (50%) | 0.8 | 0.831 |
|  | High | | 26 (53%) | 8 (50%) |  |  |
| **NK** | Low | | 26 (55%) | 6 (40%) | 0.3 | 0.502 |
|  | High | | 21 (45%) | 9 (60%) |  |  |
| **CD8^+^CD27^+^CD28^+^** | Low | | 20 (41%) | 12 (75%) | **0.018** | 0.176 |
|  | High | | 29 (59%) | 4 (25%) |  |  |
| **CD8^+^CD27^-^CD28^-^** | Low | | 26 (53%) | 6 (38%) | 0.3 | 0.502 |
|  | High | | 23 (47%) | 10 (63%) |  |  |
| **CD8^+^PD1^+^** | Low | | 22 (45%) | 10 (63%) | 0.2 | 0.502 |
|  | High | | 27 (55%) | 6 (38%) |  |  |
| **CD8^+^KLRG1^+^** | Low | | 23 (47%) | 8 (50%) | 0.8 | 0.831 |
|  | High | | 26 (53%) | 8 (50%) |  |  |
| **CD8^+^CD57^+^** | Low | | 27 (55%) | 6 (38%) | 0.2 | 0.502 |
|  | High | | 22 (45%) | 10 (63%) |  |  |
| *^1^* n (%) |  |  |  |  |  |  |
| *^2^* Pearson’s Chi-squared test |  |  |  |  |  |  |
| *^3^* Benjamini & Hochberg correction for multiple testing |  |  |  |  |  |  |

| **Myometrial infiltration** |  | | **>50%** N = 33*^1^* | **≤50%** N = 28*^1^* | **p-value***^2^* | **q-value***^3^* |
| --- | --- | --- | --- | --- | --- | --- |
| **Dendritic_cells** | Low | | 18 (55%) | 12 (43%) | 0.4 | 0.726 |
|  | High | | 15 (45%) | 16 (57%) |  |  |
| **MDSCs** | Low | | 12 (38%) | 19 (68%) | **0.019** | 0.189 |
|  | High | | 20 (63%) | 9 (32%) |  |  |
| **Tregs** | Low | | 17 (57%) | 12 (44%) | 0.4 | 0.726 |
|  | High | | 13 (43%) | 15 (56%) |  |  |
| **CD8^+^** | Low | | 16 (48%) | 12 (43%) | 0.7 | 0.865 |
|  | High | | 17 (52%) | 16 (57%) |  |  |
| **NK** | Low | | 16 (53%) | 15 (54%) | >0.9 | 0.986 |
|  | High | | 14 (47%) | 13 (46%) |  |  |
| **CD8^+^CD27^+^CD28^+^** | Low | | 16 (48%) | 14 (50%) | >0.9 | 0.986 |
|  | High | | 17 (52%) | 14 (50%) |  |  |
| **CD8^+^CD27^-^CD28^-^** | Low | | 15 (45%) | 15 (54%) | 0.5 | 0.865 |
|  | High | | 18 (55%) | 13 (46%) |  |  |
| **CD8^+^PD1^+^** | Low | | 13 (39%) | 16 (57%) | 0.2 | 0.726 |
|  | High | | 20 (61%) | 12 (43%) |  |  |
| **CD8^+^KLRG1^+^** | Low | | 14 (42%) | 16 (57%) | 0.3 | 0.726 |
|  | High | | 19 (58%) | 12 (43%) |  |  |
| **CD8^+^CD57^+^** | Low | | 16 (48%) | 15 (54%) | 0.7 | 0.865 |
|  | High | | 17 (52%) | 13 (46%) |  |  |
| *^1^* n (%) |  |  |  |  |  |  |
| *^2^* Pearson’s Chi-squared test |  |  |  |  |  |  |
| *^3^* Benjamini & Hochberg correction for multiple testing |  |  |  |  |  |  |

**Supplementary Table 4. Distribution of the peripheral immune cell profiles based on the myometrial infiltration status.**

**Supplementary Table 5. Distribution of the peripheral immune cell profiles based on the MSI status.**

| **MSI** |  | | **MSI** N = 13*^1^* | **MSS** N = 54*^1^* | **p-value***^2^* | **q-value***^3^* |
| --- | --- | --- | --- | --- | --- | --- |
| **Dendritic_cells** | Low | | 7 (54%) | 26 (48%) | 0.7 | 0.757 |
|  | High | | 6 (46%) | 28 (52%) |  |  |
| **MDSCs** | Low | | 7 (54%) | 26 (49%) | 0.8 | 0.757 |
|  | High | | 6 (46%) | 27 (51%) |  |  |
| **Tregs** | Low | | 10 (77%) | 21 (43%) | **0.029** | 0.089 |
|  | High | | 3 (23%) | 28 (57%) |  |  |
| **CD8^+^** | Low | | 9 (69%) | 24 (44%) | 0.11 | 0.155 |
|  | High | | 4 (31%) | 30 (56%) |  |  |
| **NK** | Low | | 9 (75%) | 23 (44%) | 0.055 | 0.109 |
|  | High | | 3 (25%) | 29 (56%) |  |  |
| **CD8^+^CD27^+^CD28^+^** | Low | | 3 (23%) | 30 (56%) | **0.035** | 0.089 |
|  | High | | 10 (77%) | 24 (44%) |  |  |
| **CD8^+^CD27^-^CD28^-^** | Low | | 10 (77%) | 23 (43%) | **0.026** | 0.089 |
|  | High | | 3 (23%) | 31 (57%) |  |  |
| **CD8^+^PD1^+^** | Low | | 9 (69%) | 24 (44%) | 0.11 | 0.155 |
|  | High | | 4 (31%) | 30 (56%) |  |  |
| **CD8^+^KLRG1^+^** | Low | | 7 (54%) | 26 (48%) | 0.7 | 0.757 |
|  | High | | 6 (46%) | 28 (52%) |  |  |
| **CD8^+^CD57^+^** | Low | | 10 (77%) | 23 (43%) | **0.026** | 0.089 |
|  | High | | 3 (23%) | 31 (57%) |  |  |
| *^1^* n (%) |  |  |  |  |  |  |
| *^2^* Pearson’s Chi-squared test |  |  |  |  |  |  |
| *^3^* Benjamini & Hochberg correction for multiple testing |  |  |  |  |  |  |

**Supplementary Table 6. Distribution of the peripheral immune cell profiles based on the P53 status.**

| **P53** |  | | **MUT** N = 16*^1^* | **WT** N = 44*^1^* | **p-value***^2^* | **q-value***^3^* |
| --- | --- | --- | --- | --- | --- | --- |
| **Dendritic_cells** | Low | | 9 (56%) | 20 (45%) | 0.5 | 0.656 |
|  | High | | 7 (44%) | 24 (55%) |  |  |
| **MDSCs** | Low | | 10 (63%) | 22 (50%) | 0.4 | 0.656 |
|  | High | | 6 (38%) | 22 (50%) |  |  |
| **Tregs** | Low | | 4 (27%) | 24 (59%) | **0.035** | 0.347 |
|  | High | | 11 (73%) | 17 (41%) |  |  |
| **CD8^+^** | Low | | 6 (38%) | 23 (52%) | 0.3 | 0.656 |
|  | High | | 10 (63%) | 21 (48%) |  |  |
| **NK** | Low | | 7 (44%) | 23 (55%) | 0.5 | 0.656 |
|  | High | | 9 (56%) | 19 (45%) |  |  |
| **CD8^+^CD27^+^CD28^+^** | Low | | 9 (56%) | 20 (45%) | 0.5 | 0.656 |
|  | High | | 7 (44%) | 24 (55%) |  |  |
| **CD8^+^CD27^-^CD28^-^** | Low | | 7 (44%) | 22 (50%) | 0.7 | 0.835 |
|  | High | | 9 (56%) | 22 (50%) |  |  |
| **CD8^+^PD1^+^** | Low | | 8 (50%) | 20 (45%) | 0.8 | 0.839 |
|  | High | | 8 (50%) | 24 (55%) |  |  |
| **CD8^+^KLRG1^+^** | Low | | 8 (50%) | 21 (48%) | 0.9 | 0.876 |
|  | High | | 8 (50%) | 23 (52%) |  |  |
| **CD8^+^CD57^+^** | Low | | 6 (38%) | 25 (57%) | 0.2 | 0.656 |
|  | High | | 10 (63%) | 19 (43%) |  |  |
| *^1^* n (%) |  |  |  |  |  |  |
| *^2^* Pearson’s Chi-squared test |  |  |  |  |  |  |
| *^3^* Benjamini & Hochberg correction for multiple testing |  |  |  |  |  |  |

**Supplementary Table 7. Distribution of the peripheral immune cell profiles based on the therapy response in advanced endometrial cancer patients.**

|  | **Relapse** | **Non-relapse** N = 16*^1^* | **Relapse** N = 15*^1^* | **p-value***^2^* | **q-value***^3^* |
| --- | --- | --- | --- | --- | --- |
| **Dendritic_cells** | Low | 8 (50%) | 7 (47%) | 0.9 | >0.999 |
|  | High | 8 (50%) | 8 (53%) |  |  |
| **MDSCs** | Low | 8 (50%) | 7 (50%) | >0.9 | >0.999 |
|  | High | 8 (50%) | 7 (50%) |  |  |
| **Tregs** | Low | 8 (57%) | 5 (42%) | 0.4 | 0.616 |
|  | High | 6 (43%) | 7 (58%) |  |  |
| **CD8^+^** | Low | 11 (69%) | 4 (27%) | **0.019** | 0.064 |
|  | High | 5 (31%) | 11 (73%) |  |  |
| **NK** | Low | 7 (50%) | 7 (50%) | >0.9 | >0.999 |
|  | High | 7 (50%) | 7 (50%) |  |  |
| **CD8^+^CD27^+^CD28^+^** | Low | 5 (31%) | 10 (67%) | **0.049** | 0.122 |
|  | High | 11 (69%) | 5 (33%) |  |  |
| **CD8^+^CD27^-^CD28^-^** | Low | 12 (75%) | 3 (20%) | **0.002** | **0.022** |
|  | High | 4 (25%) | 12 (80%) |  |  |
| **CD8^+^PD1^+^** | Low | 6 (38%) | 9 (60%) | 0.2 | 0.421 |
|  | High | 10 (63%) | 6 (40%) |  |  |
| **CD8^+^KLRG1^+^** | Low | 9 (56%) | 6 (40%) | 0.4 | 0.609 |
|  | High | 7 (44%) | 9 (60%) |  |  |
| **CD8^+^CD57^+^** | Low | 11 (69%) | 4 (27%) | **0.019** | 0.086 |
|  | High | 5 (31%) | 11 (73%) |  |  |
| *^1^* n (%) |  | | | |  |
| *^2^* Pearson’s Chi-squared test |  | | | |  |
| *^3^* Benjamini & Hochberg correction for multiple testing |  | | | |  |

**Supplementary Table 8. Univariate Cox Analysis of immune cell subsets in advanced EC.**

| **Cell Type** | **N** | **Events** | **HR** | **CI_Lower** | **CI_Upper** | **p_value** | **q_value** |
| --- | --- | --- | --- | --- | --- | --- | --- |
| **Dendritic_cells** | 31 | 15 | 1.006 | 0.766 | 1.320 | 0.967 | 0.967 |
| **MDSCs** | 30 | 14 | 0.976 | 0.814 | 1.171 | 0.797 | 0.886 |
| **NK** | 28 | 14 | 0.986 | 0.923 | 1.055 | 0.690 | 0.862 |
| **CD8^+^CD27^+^CD28^+^** | 31 | 15 | 0.963 | 0.933 | 0.995 | **0.022** | **0.044** |
| **CD8^+^CD27^-^CD28^-^** | 31 | 15 | 1.043 | 1.015 | 1.072 | **0.003** | **0.018** |
| **CD8^+^** | 31 | 15 | 1.057 | 1.016 | 1.099 | **0.006** | **0.021** |
| **CD8^+^KLRG1^+^** | 31 | 15 | 1.017 | 0.993 | 1.042 | 0.166 | 0.276 |
| **CD8^+^CD57^+^** | 31 | 15 | 1.063 | 1.020 | 1.108 | **0.004** | **0.018** |
| **CD8^+^PD1^+^** | 31 | 15 | 0.984 | 0.950 | 1.020 | 0.388 | 0.554 |
| **Low_mature_high_senescent_CD8^+^** | 31 | 15 | 1.014 | 1.003 | 1.025 | **0.015** | **0.037** |

**Supplementary Table 9.** **Exploratory Multivariable Cox models adjusted for age, histology, grade and myometrial infiltration in advanced EC.**

|  |  |  | |  |  |  |  |  |
| --- | --- | --- | --- | --- | --- | --- | --- | --- |
| **Cell_Type** | **N** | **Events** | **HR** | | **CI_Lower** | **CI_Upper** | **p_value** | **q_value** |
| **CD8^+^** | 23 | 14 | 1.134 | | 1.049 | 1.227 | **0.002** | **0.005** |
| **CD8^+^CD27^-^CD28^-^** | 23 | 14 | 1.039 | | 1.009 | 1.071 | **0.011** | **0.011** |
| **CD8^+^CD57^+^** | 23 | 14 | 1.069 | | 1.021 | 1.119 | **0.004** | **0.006** |

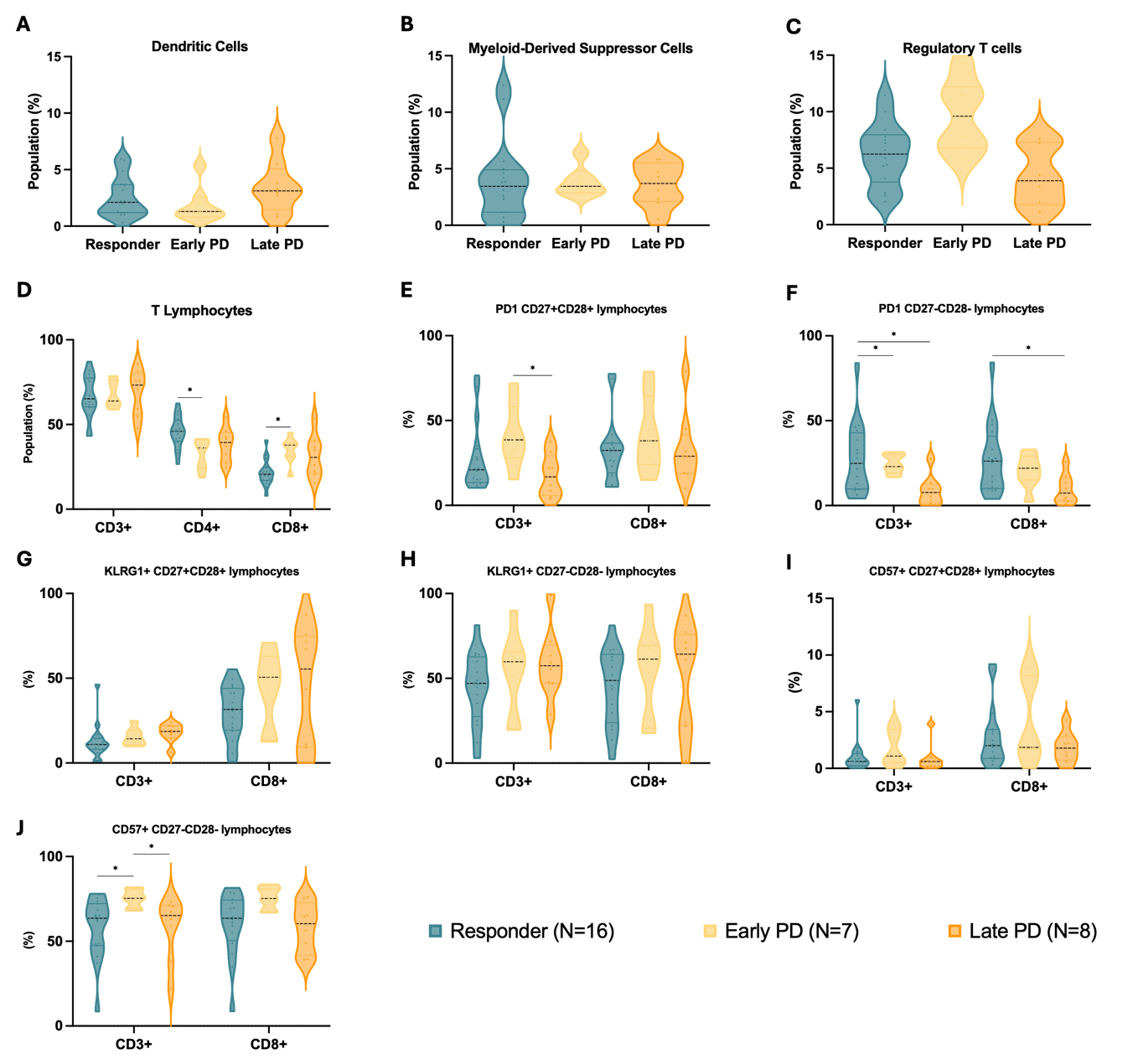

**Supplementary Figure 4. Immune cell profiles in advanced EC patients. A.** Dendritic levels across Responders, Early PD and Late PD. **B.** MDSCs. **C.** Regulatory T cells. **D.** T lymphocytes. **E.** PD1^+^CD27^+^CD28^+^ cells. **F.** PD1^+^CD27^-^CD28^-^ cells. **G.** KLRG1^+^CD27^+^CD28^+^ cells. **H.** KLRG1^+^CD27^-^CD28^-^ cells. **I.** CD57^+^CD27^+^CD28^+^ cells. **J.** CD57^+^CD27^-^CD28^-^ cells. Data are presented as mean ± standard deviation. **p* < 0.05, ***p* < 0.01, ****p* < 0.001, Kruskal-Wallis test with Dunn's post-hoc analysis.
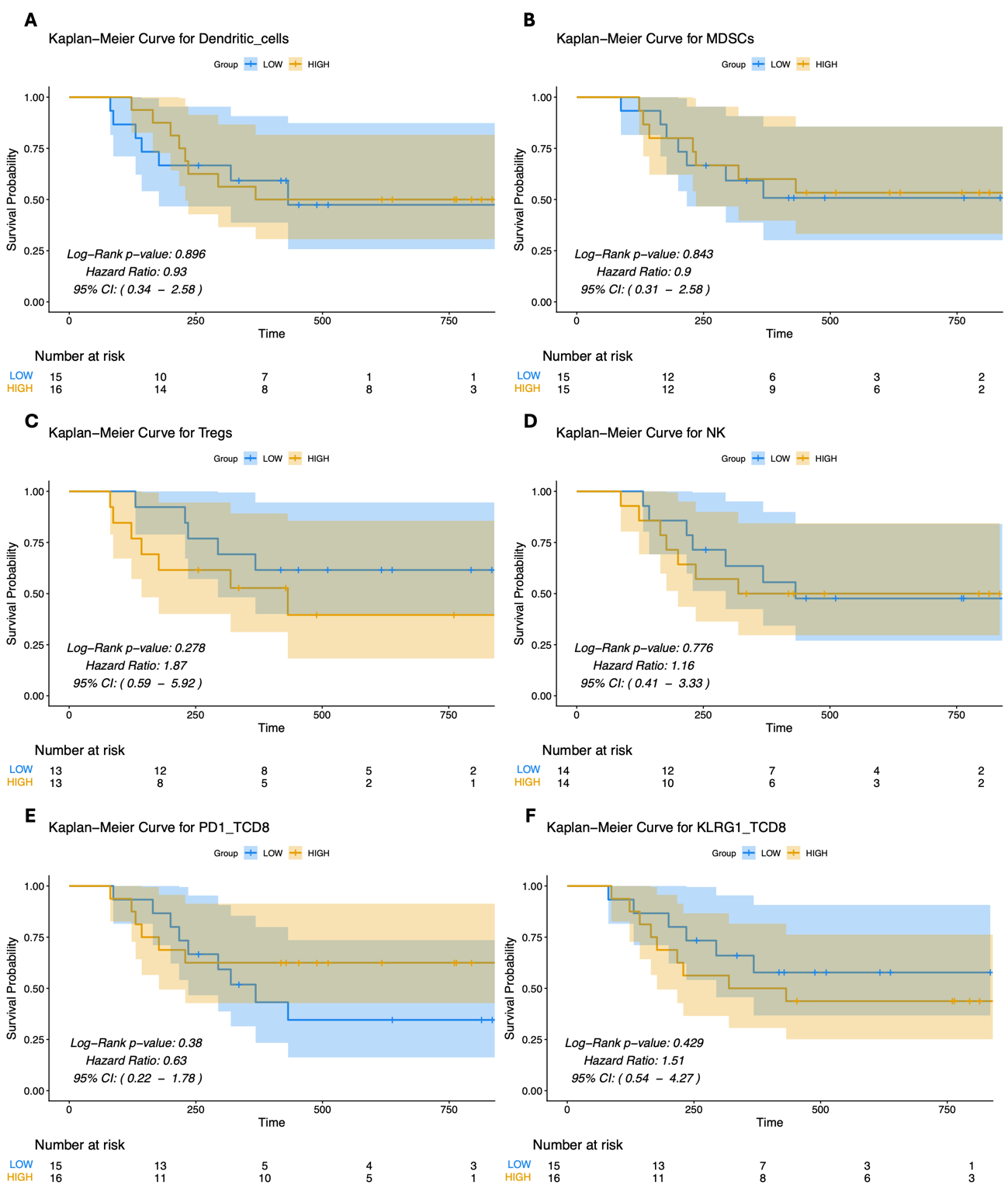

**Supplementary Figure 5. Prognostic Impact of Immune Cell Subtypes on PFS in EC Patients. A**. Kaplan-Meier plot of PFS among patients with high levels of dendritic cells. **B.** Similarly, for patients with high levels MDSCs. **C.** Kaplan-Meier plot for Tregs**. D.** For NK cells. **E.** For PD-1^+^ immune cells. **F.** For KLRG1^+^ lymphocytes. **G.** Kaplan-Meier plot for CD57^+^ senescent T cells. Statistical comparisons were performed using Log-Rank tests.

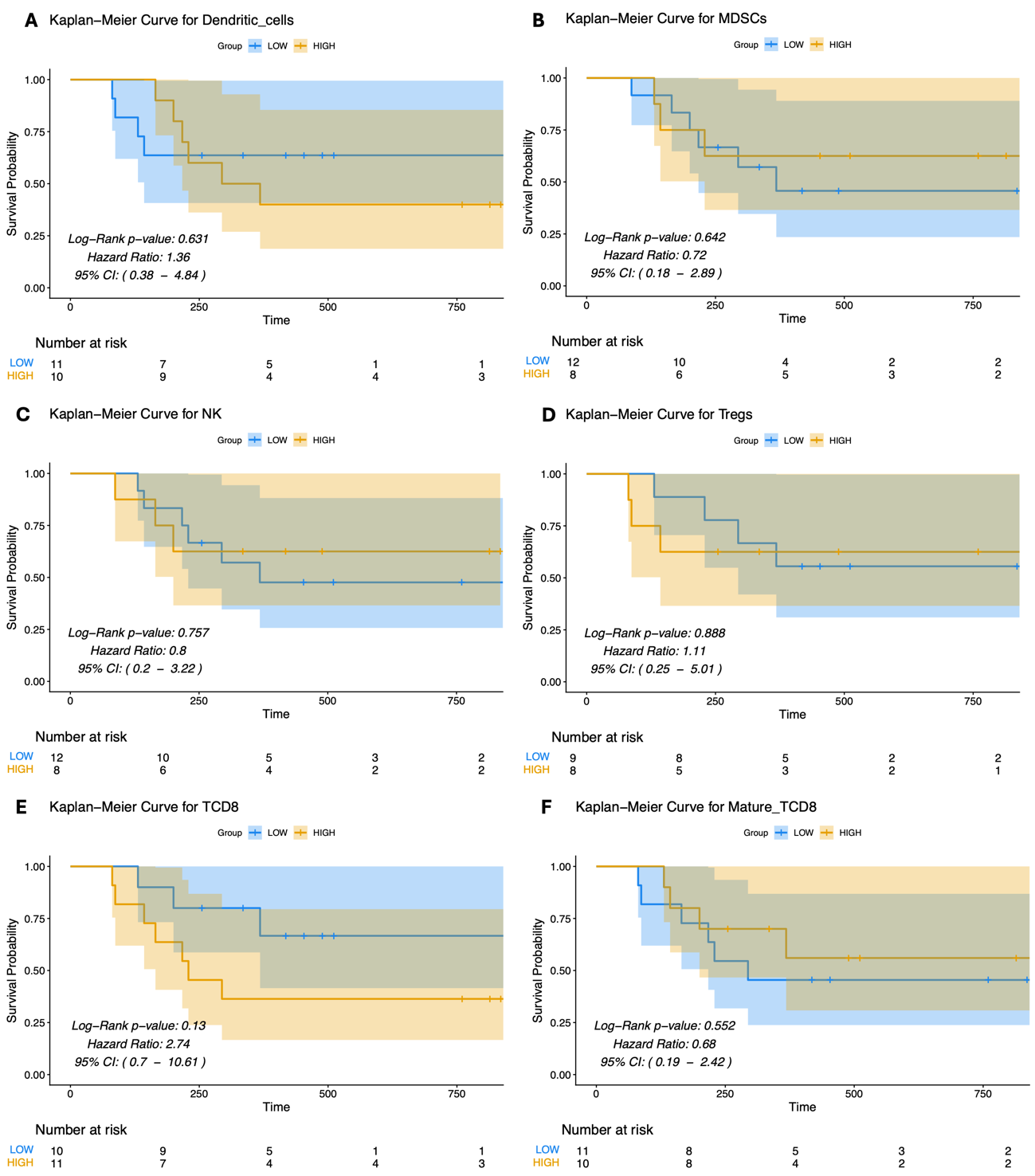

**Supplementary Figure 6. Prognostic Impact of Immune Cell Subtypes on PFS in Advanced EC Patients treated with chemotherapy**. Kaplan-Meier plot of PFS among patients according to **A.** their levels of dendritic cells, **B.** MDSCs, **C.** NK**, D.** Regulatory T cells, **E.** CD8^+^ T cells, **F.** mature CD8^+^ T cells, **G.** for patients with low levels of mature and high levels of exhausted CD8^+^ T cells, **H.** KLRG1^+^ T cells, **I.** PD-1^+^CD8^+^T cells and **J.** CD57^+^CD8^+^ T cells. Statistical comparisons were performed using log-rank tests.

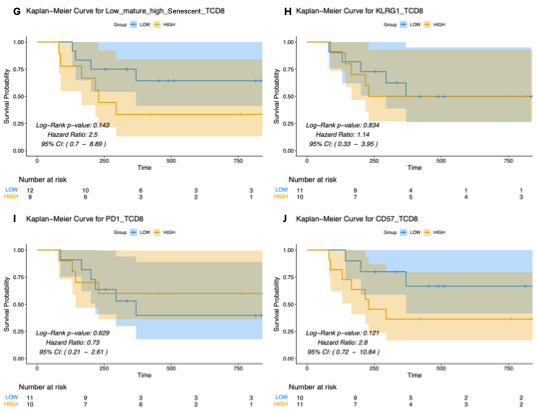

**Supplementary Figure 6 (cont)**.

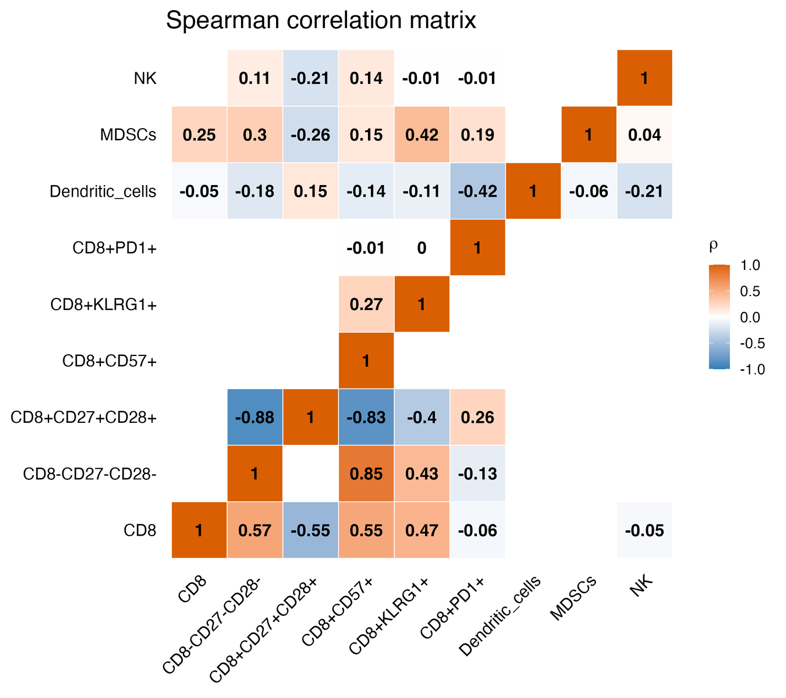
**Supplementary Figure 7. Spearman correlation heatmap of the analyzed immune markers.**

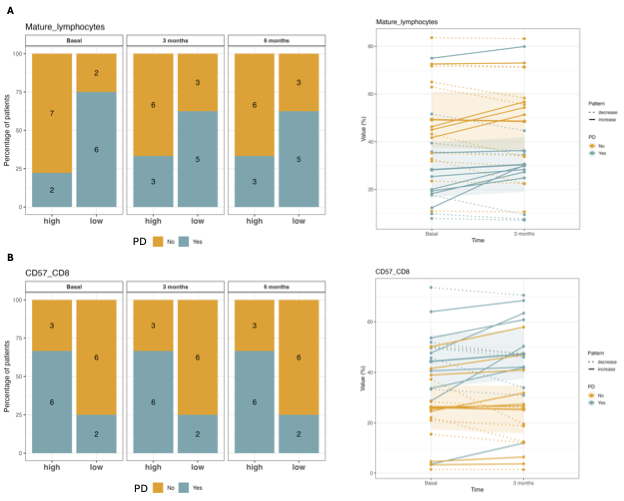
**Supplementary Figure 8. Longitudinal analysis of peripheral blood immune cell subpopulations with advanced EC. A.** Mature CD8^+^ lymphocytes. **B.** CD8^+^CD57^+^ lymphocytes. Barplots (left panels) show the percentage of patients with high and low values of each population at baseline, 3 months, and 6 months. Paired-line plots (right panels) illustrate individual trajectories for each cell subset, stratified by disease progression (PD: Yes/No).
